# A>G substitutions on a heavy chain of mitochondrial genome marks an increased level of aerobic metabolism in warm versus cold vertebrates

**DOI:** 10.1101/2020.07.25.221184

**Authors:** Alina G. Mikhailova, Dmitrii Iliushchenko, Victor Shamansky, Alina A. Mikhailova, Kristina Ushakova, Evgenii Tretyakov, Sergey Oreshkov, Dmitry Knorre, Leonard Polishchuk, Dylan Lawless, Aleksandr Kuzmin, Stepan Denisov, Ivan Kozenkov, Ilya Mazunin, Wolfram Kunz, Masashi Tanaka, Vsevolod Makeev, Rita Castilho, Valerian Yurov, Alexander Kuptsov, Jacques Fellay, Konstantin Khrapko, Konstantin Gunbin, Konstantin Popadin

## Abstract

The variation in the mutational spectrum of the mitochondrial genome (mtDNA) among species is not well understood. Recently, we demonstrated an increase in A>G substitutions on a heavy chain (hereafter A_H_>G_H_) of mtDNA in aged mammals, interpreting it as a hallmark of age-related oxidative damage. In this study, we hypothesized that the occurrence of A_H_>G_H_ substitutions may depend on the level of aerobic metabolism, which can be inferred from an organism’s body temperature. To test this hypothesis, we used body temperature in endotherms and environmental temperature in ectotherms as proxies for metabolic rate and reconstructed mtDNA mutational spectra for 1350 vertebrate species. Our results showed that temperature was associated with increased rates of A_H_>G_H_ and asymmetry of A_H_>G_H_ in different species of ray-finned fishes and within geographically distinct clades of European anchovy. Analysis of nucleotide composition in the most neutral synonymous sites of fishes revealed that warm-water species were expectedly more A-poor and G-rich compared to cold-water species. Finally, we extended our analyses to all vertebrates and observed higher A_H_>G_H_ and increased asymmetry of A_H_>G_H_ in warm-blooded (mammals and birds) compared to cold-blooded (Actinopterygii, amphibia, reptilia) vertebrate classes. Overall, our findings suggest that temperature, through its influence on metabolism and oxidative damage, shapes the mutational properties and nucleotide content of the mtDNA in all vertebrates.

## INTRODUCTION

A high mitochondrial mutation rate provides a rich source of variants that are extensively used in tracing the history of species, populations, organisms within populations, and, more recently - cells in tissues (Ludwig et al.2019). Here we propose to extend the utility of neutral mtDNA polymorphic data by deriving species-specific mutational spectra and correlating them with life-history traits. We have shown recently that mammalian mtDNA mutational spectrum, and precisely A_H_>G_H_ substitution (hereafter index _H_ marks mtDNA heavy chain annotation, Methods), is associated with mammalian lifespan (Mikhailova et al. 2022), most likely through age-related oxidative damage. Here, we hypothesize that the mtDNA mutational spectrum can also be sensitive to oxidative damage through the species-specific rate of aerobic metabolism. Since the level of metabolism depends strongly on temperature (J. F. Gillooly et al. 2001), we decided to analyze the mtDNA mutational spectrum across different ectothermic and endothermic vertebrates. Effects of temperature on mutation rate have been intensively studied for decades (Timoféeff-Ressovsky 1936; Lindgren 1972; James F. Gillooly et al. 2005; Chu et al. 2018; Belfield et al. 2021; Matsuba et al. 2013) however, temperature-specific mutational signature - i.e. the effect of temperature on changes in mutational spectrum became a focus of first studies just recently (Waldvogel and Pfenninger 2021; Chu et al. 2018) where mtDNA was not under the focus.

## RESULTS

### 1. Species-specific mtDNA mutational spectrum of fishes is associated with ambient water temperature by means of increased asymmetry of A_H_>G_H_

In order to investigate the potential effect of temperature on mtDNA mutagenesis, we focused on fishes (Actinopterygii N=180; Chondrichthyes N=8), which are ectothermic animals that inhabit a range of ambient water temperatures. Using within-species synonymous mtDNA polymorphisms, we reconstructed a 12-component mutational spectrum for each of the 188 fish species (see Methods and Supplementary File 1). The average mutational spectrum displayed a clear excess of transitions, with C_H_>T_H_ being the most common and A_H_>G_H_ being the second most common (Fig. 1A). This pattern closely resembled the spectra seen in mammals (Mikhailova et al. 2022) and human cancers (Yuan et al. 2020; Ju et al. 2014). Next, we collected data on the mean annual water temperatures for each fish species (Methods) and analyzed its relationship with the 12 types of substitutions in the mtDNA mutational spectrum.

**FIG1.**
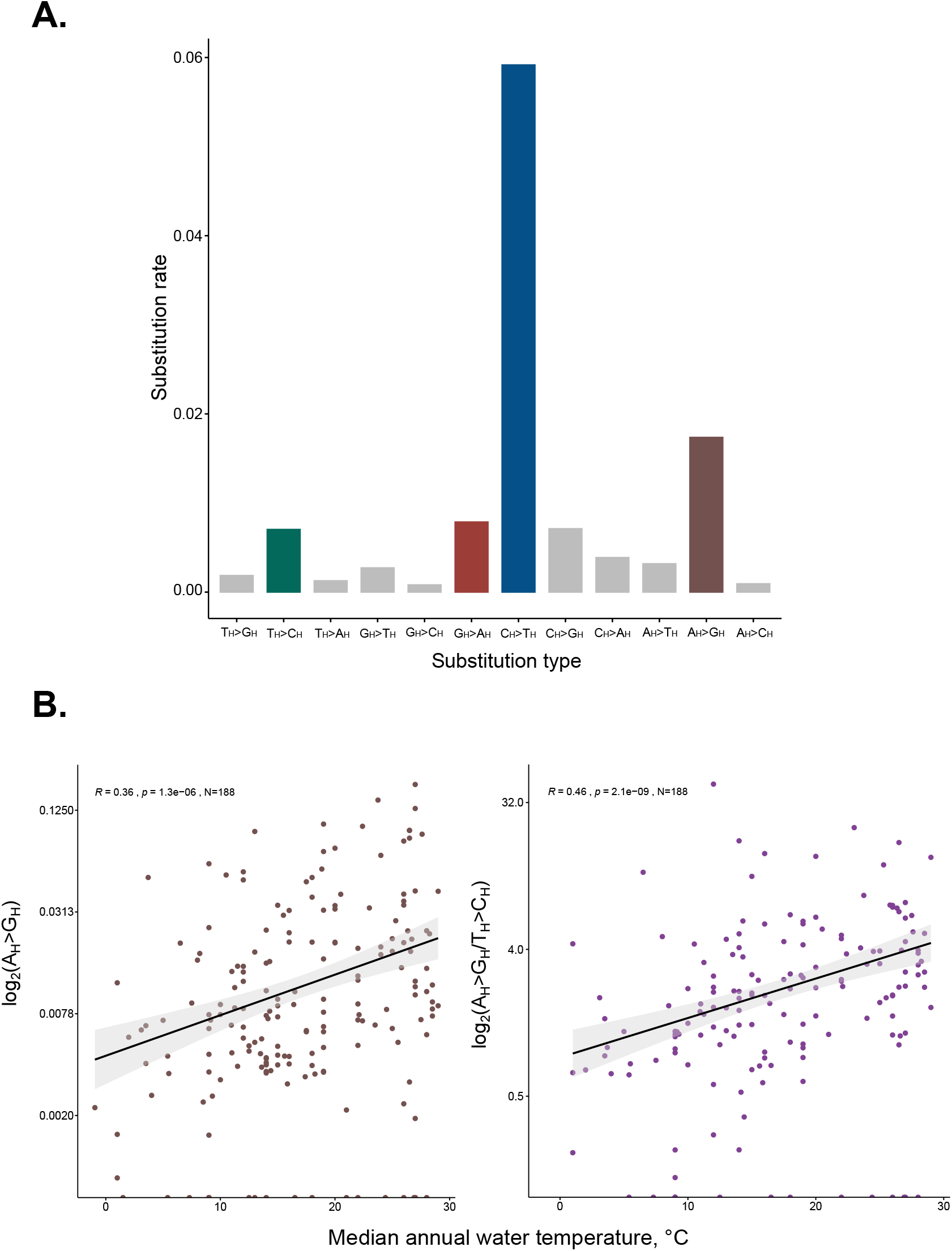
mtDNA mutational spectrum of Actinopterygii species (mtDNA heavy chain notation). (A) average mutational spectrum of all analyzed Actinopterygii species; (B) species-specific mutational spectrum is associated with water temperature. Left panel: positive correlation of A_H_>G_H_ with temperature (Spearman’s rho = −0.36, p-value = 3.3e-05, N = 128); middle panel: negative correlation of T_H_>C_H_ with temperature (Spearman’s rho = 0.26, p-value = 0.0025, N = 128); right panel: positive correlation of log2(Ah>Gh/Th>Ch) with temperature (Spearman’s rho = 0.44, p-value = 3.1e-07, N = 123)

We found that two types of substitutions, A_H_>G_H_ and C_H_>T_H_, had a significant positive correlation with temperature (Fig. 1B, Supplementary Mat. 1a). To determine whether the asymmetry of these substitutions also increased with temperature, we examined corresponding ratios A_H_>G_H_/T_H_>C_H_ and C_H_>T_H_/G_H_>A_H_. The logic behind the asymmetry analysis is so that, for example, the ratio A_H_>G_H_/T_H_>C_H_ represents an asymmetry of A_H_>G_H_ substitutions. A>G and T>C are complementary equivalents: a substitution annotated as A>G on the heavy chain (A_H_>G_H_) might have originated as T>C on the light chain (T_L_>C_L_), and vice versa. If the probability of A mutating into G is the same on both the heavy (A_H_>G_H_) and light (A_L_>G_L_ = T_H_>C_H_) chains, we would expect to see symmetry, with the rate of A_H_>G_H_ equal to the rate of T_H_>C_H_ (both rates are normalized by the frequency of ancestral nucleotides). However, we observed that both asymmetries are strong enough: A_H_>G_H_ was higher than T_H_>C_H_ (A_H_>G_H_/ T_H_>C_H_ > 1, see figure 1A), and C_H_>T_H_ was higher than G_H_>A_H_ (C_H_>T_H_/G_H_>A_H_> 1, see figure 1A) indicating mutagens that predominantly induced A>G and C>T on the heavy chain (Supplementary Fig. 1.1). Next, we observed that A_H_>G_H_ asymmetry, but not C_H_>T_H_ asymmetry, demonstrated a strong positive correlation with temperature (Fig. 1B, Supplementary Mat. 1a,1b). The increased A_H_>G_H_ asymmetry in warm-versus cold-water species shows that this mutagen is sensitive to temperature - i.e. induces proportionally more A>G on heavy versus light chain with an increase of temperature. This trend is robust to phylogenetic inertia (Supplementary Mat. 1c), and remains qualitatively similar when we analyze family-specific data and when we split the entire dataset into several subgroups based on temperature (Supplementary Fig. 1.2).

Previous research has shown that A_H_>G_H_ in mammalian species is positively correlated with generation length (Mikhailova et al. 2022). To examine if a similar relationship exists in fish, we used an analogous metric - the time of maturation estimated as the age at which 50% of a cohort spawn for the first time. We conducted a set of rank correlations and multiple linear regression analyses to investigate the effect of temperature and the time of maturation on the mtDNA mutational spectrum, specifically the A_H_>G_H_, A_H_>G_H_ asymmetry, C_H_>T_H_, C_H_>T_H_ asymmetry (Supplementary Mat. 1d, 1e, 1f, 1g). First of all, we observed a negative correlation between the temperature and the time of maturation (Supplementary Mat. 1d) suggesting that warm-water species tend to reach maturation faster. Second, neither of the key four metrics of the mutational spectrum showed correlations with the time of maturation (Supplementary Mat. 1e, 1f, 1g). Overall, we conclude that the fish mtDNA mutational spectrum, particularly the A_H_>G_H_ and A_H_>G_H_ asymmetry, and weakly C_H_>T_H_ are associated with the temperature only.

Previous research has shown that A_H_>G_H_ in the mtDNA of mammalian species is positively correlated with generation length (Mikhailova et al. 2022). To investigate whether a similar relationship exists in fish, we used an analogous metric: the time of maturation, defined as the age at which 50% of a cohort reaches sexual maturity. We conducted a series of rank correlations and multiple linear regression analyses to examine the effect of temperature and the time of maturation on the mtDNA mutational spectrum, specifically the A_H_>G_H_, AH>GH asymmetry, CH>TH, CH>TH asymmetry (Supplementary Mat. 1d, 1e, 1f, 1g). We observed a negative correlation between temperature and time of maturation (Supplementary Mat. 1d), suggesting that warm-water species tend to reach sexual maturity faster. However, none of the key four metrics of the mutational spectrum showed correlations with the time of maturation (Supplementary Mat. 1e, 1f, 1g). Overall, our results suggest that the fish mtDNA mutational spectrum, particularly the A_H_>G_H_ and A_H_>G_H_ asymmetry, and weakly C_H_>T_H_, are associated with the temperature only.

### 2. Tropical clade of European anchovy shows also an increased asymmetry of A_H_>G_H_

Widely distributed species with genetic clinal variation over geographical ranges is an ideal model to test our findings on the within-species scale. Intensive studies of the European anchovy (*Engraulis encrasicolus*) uncovered the existence of two mtDNA clades, clonally arranged along the eastern Atlantic Ocean: clade A is preferentially distributed in the tropics, while clade B has an anti-tropical distribution, with frequencies decreasing towards the tropics (Silva et al. 2014). Analysing a fragment of cytochrome B for 285 organisms of clade A and 160 organisms of clade B (data generously provided by Dr. Rita Castilho), we reconstructed their mutational spectra: for each clade, we reconstructed a consensus and based on all synonymous deviations from consensus we derived the mutational spectra (Methods). We observed that both A_H_>G_H_ and A_H_>G_H_ asymmetry (A_H_>G_H_/T_H_>C_H_) were higher in tropical clade A (Fig. 2); frequencies of all other types of substitutions were similar. The observed pattern is completely in line with our previous findings, showing an increased A_H_>G_H_ and increased A_H_>G_H_ asymmetry in warm-water species (Fig. 1B).

**FIG2.**
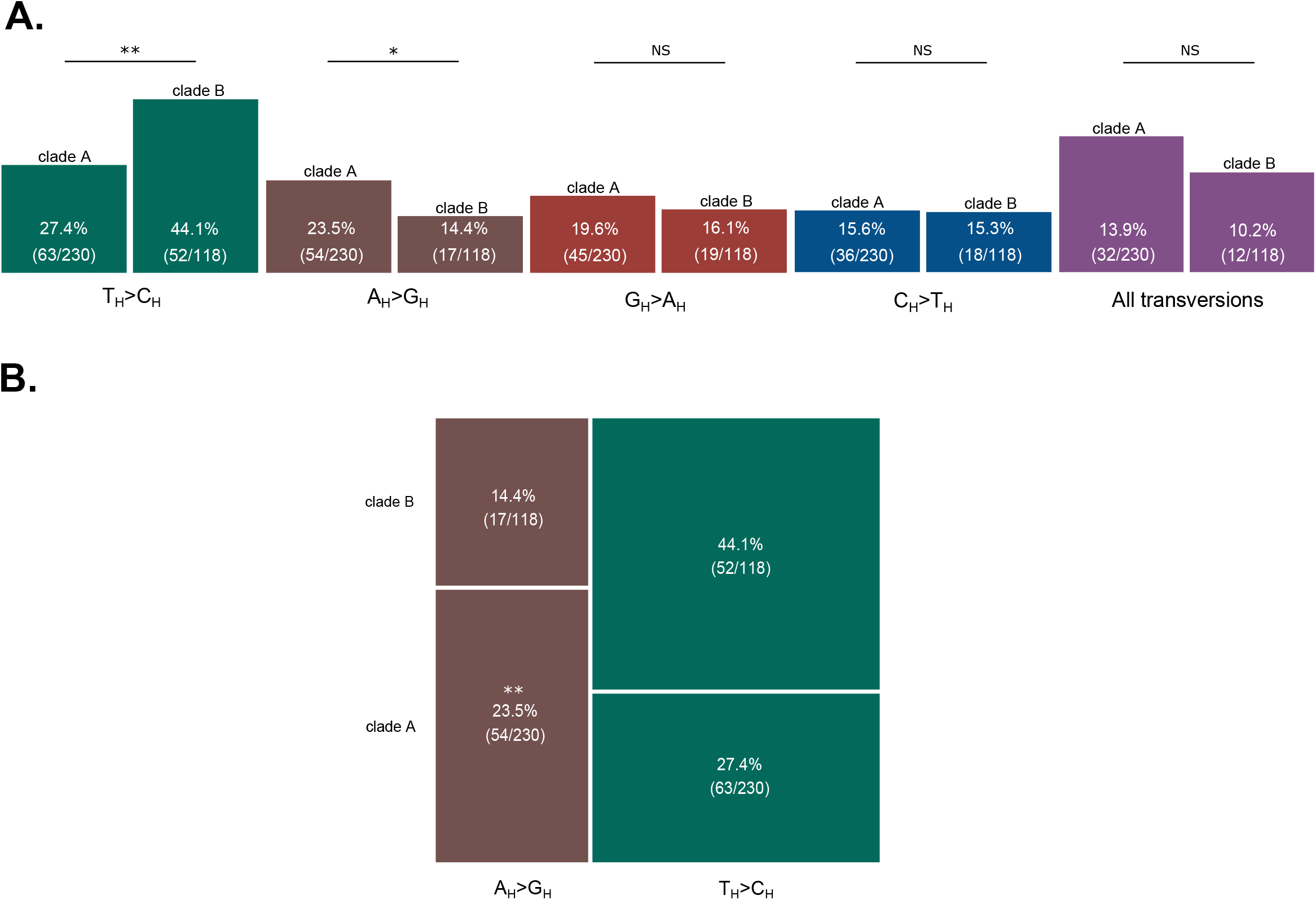
mtDNA mutational spectrum of European anchovy clades. (A) Synonymous substitutions, reconstructed for clades A and B were split into five groups: four types of transitions and one group of transversions (all transversions were merged together due to their rarity). More tropically distributed clade A shows an excess of A_H_>G_H_ (Fisher’s odds ratio = 1.81, p-value = 0.049) and a deficit of T_H_>C_H_ (Fisher’s odds ratio = 0.48, p-value = 0.0025). Frequencies of all other types of substitutions were similar (all p-values > 0.5). (B) Analysis of only two fluctuating types of transitions: A_H_>G_H_ versus T_H_>C_H_ between clades A and B emphasizes an excess of A_H_>G_H_ as compared to T_H_>C_H_ in tropical clade A (Fisher’s odds ratio = 2.6, p = 0.005). One asterisk denotes p value < 0.05, two asterisks mark p value < 0.01. Two asterisks mark p value < 0.01

The mutational spectrum, with time, can affect nucleotide composition, especially in synonymous positions. Assuming that consensus sequences of clades A and B reflect the ancestral states, we compared their nucleotide content in synonymous positions. Totally we observed 21 synonymous differences between consensuses of clades A and B and, interestingly, nucleotide composition in these sites was completely in line with the expected mutational bias: tropical clade A versus B is more A poor (3 versus 10), G rich (9 versus 4), T rich (7 versus 1) and C poor (2 versus 6) (Supplementary Mat. 2a). All these changes might result from the clade-specific mutational spectra: an increased A_H_>G_H_ and A_H_>G_H_ asymmetry in tropical clade A (Fig 2). It is important to emphasize that observed divergence in the consensus sequences between clades A and B makes the difference in the mutational spectra even more pronounced and robust. For example, clade A demonstrates an excess of substitutions A_H_>G_H_ despite the deficit of A_H_ among synonymous sites of the consensus. Altogether we observed an increased asymmetry of A_H_>G_H_ in the tropical clade of European anchovy as well as a line of evidence that the mutational bias affects the neutral nucleotide composition of the sequences.

### 3. The mtDNA mutational spectrum of fish impacts their neutral nucleotide composition: warm water species tend to be AC-poor and GT-rich

The nucleotide composition can be influenced by the mutational spectrum: if mutational bias is stronger than selection and the analyzed species are close to nucleotide compositional equilibrium, the mutational spectrum in warm-water fishes ultimately leads to a decrease in A_H_ and an increase in G_H_ over the long term.

To determine how close fishes are to their nucleotide compositional equilibrium, we compared the expected and observed nucleotide compositions in the most neutral, four-fold degenerate positions (see Methods). First, we calculated the typical mutational spectra for cold- and warm-water fishes using the coldest (N=19, temp <= 8.18 °C) and warmest (N= 23, temp >= 27°C) deciles of the analyzed species. As expected, warm-water fishes exhibited increased A_H_>G_H_ and A_H_>G_H_ asymmetry (Supplementary Mat. 3a). Second, using computer simulations and analytical solutions, we derived the expected neutral nucleotide compositions for cold and warm-water fishes based on these mutational spectra (both approaches converged on identical results, see Supplementary Fig. 3.1). This allowed us to answer the question of what the nucleotide composition would be in the long term if the only factor affecting molecular evolution was mutagenesis (i.e., the specific mutational spectrum). Third, we obtained the observed nucleotide composition by analyzing the synonymous, four-fold degenerate nucleotide content in 12 (all except ND6) protein-coding mtDNA genes from the coldest and warmest deciles, and compared it to the expected composition (Fig. 3A). We found that the observed nucleotide composition was similar to the expected composition, indicating that the analyzed species are close to compositional equilibrium. Additionally, we observed that warm-water species tend to be closer to the expected equilibrium (as indicated by the horizontal dotted lines on Fig. 3A) compared to cold-water species, possibly due to their increased mutational rate and faster approach to equilibrium.

**FIG3.**
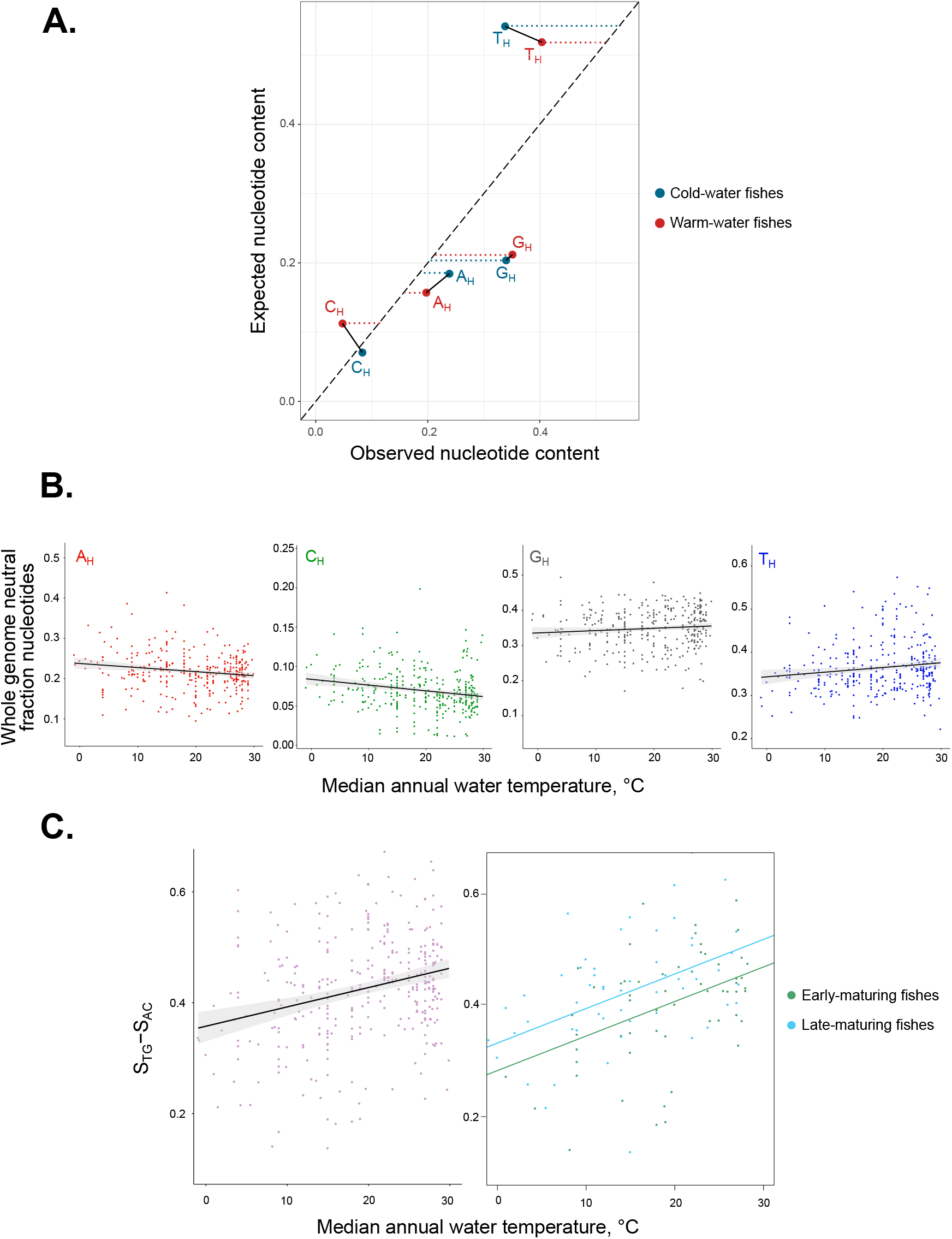
mtDNA mutational spectrum of Actinopterygii (mtDNA heavy chain notation). (A) observed and expected nucleotide compositional equilibrium of warm- and cold-water fishes. Expected nucleotide composition has been estimated based on computational simulations and analytical solutions (Supplementary Figure 3.1) (B) whole-genome neutral nucleotide content is associated with ambient temperature (A_H_: Spearman’s rho = 0.117, p-value = 0.03301; T_H_: Spearman’s rho = −0.153, p-value = 0.00537; G_H_: Spearman’s rho = −0.249, p-value = 4.711e-06; C_H_: Spearman’s rho = 0.132, p-value = 0.01655, N = 123). (C) S_TG_-S_AC_ is sensitive to both temperature and the time of maturation, temperature being the strongest factor. Left panel shows a positive relationship between S_TG_-S_AC_ and temperature for 333 species (statistical details are in supplementary Mat. 3d and 3c). Right panel demonstrates similar positive slopes between S_TG_-S_AC_ and temperature for early and late maturing fishes (N = 131), where late-maturing ones have an increased intercept (statistical details are in supplementary Mat. 3e and 3f).

Given that the mtDNA synonymous nucleotide composition of fish is close to neutral equilibrium and there is no strong selection on four-fold degenerate synonymous positions in mtDNA, we expect to observe a correlation between nucleotide content and ambient temperature. Upon analyzing the synonymous, four-fold degenerate nucleotide content in 12 (all except ND6) protein-coding genes of fish, we observed four trends: a decrease in A_H_ and C_H_ and an increase in G_H_ and T_H_ in warm water versus cold water species (Fig. 3B, Supplementary Mat. 3b). It is important to note that all of these changes reflect the consequences of increased A_H_>G_H_ and C_H_>T_H_ ratios in warm water fish. To clarify these four regressions we combined them into GHAHskew and THCHskew and analyzed their relationship with temperature, time of maturation, and both of them in a set of models (Supplementary Mat. 3c-i).

To highlight this, we calculated the sum of the fractions of gainer nucleotides - T_H_ and G_H_ (S_TG_) and subtracted from it the sum of loser nucleotides - A_H_ and C_H_ (S_AC_), creating a single metric S_TG_-S_AC_(Methods). Regressing S_TG_-S_AC_ on temperature resulted in a strong, expected positive relationship (Fig. 3C left panel, Supplementary Mat. 3d), which is robust to phylogenetic inertia (Supplementary Mat. 3e) and remains significant in two separate groups of fish: short-lived and long-lived (Supplementary Fig. 3.2).

Considering the increased A_H_>G_H_ ratio in long-lived versus short-lived mammals (Mikhailova et al. 2022), we examined the potential association of S_TG_-S_AC_ with both temperature and the time of maturation in fish. The result of the multiple linear regression analysis is as follows:

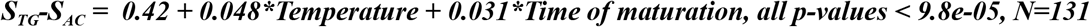

We see that both factors affect the S_TG_-S_AC_ positively, but the temperature has a higher impact on the S_TG_-S_AC_ as compared to the time of maturation (all coefficients are standardized, Fig. 3C right panel, see also Supplementary Mat. 3f). Both temperature and the time of maturation marginally significantly affect S_TG_-S_AC_ taking into account the phylogenetic inertia (Supplementary Mat. 3g). Of note, the effect of the time of maturation was not significant in our analysis of the 12-component mutational spectrum (chapter 1). Altogether, we conclude that the mtDNA nucleotide content of ray-finned fishes is affected by two factors: a strong temperature-dependent mutagen and a two-times weaker longevity-associated mutagen.

We found that both temperature and the time of maturation have a positive effect on S_TG_-S_AC_, but the temperature has a greater impact compared to the time of maturation (all coefficients are standardized, Fig. 3C right panel, see also Supplementary Mat. 3e). Both temperature and the time of maturation have a marginally significant effect on S_TG_-S_AC_ when considering phylogenetic inertia (Supplementary Mat. 3f-i). In conclusion, we find that the mtDNA nucleotide content of ray-finned fishes is influenced by two factors: a strong, temperature-dependent mutagen and a weaker, longevity-associated mutagen

### 4. The mtDNA mutational spectrum is temperature sensitive in all classes of vertebrates

The temperature sensitivity of mtDNA mutational spectra in fish species (Fig 1-3) suggests that this phenomenon may be universal. To test this, we compared the mtDNA mutational spectra of five classes of vertebrates, grouped into cold-blooded (Actinopterygii, Amphibia, Reptilia) and warm-blooded (Mammalia, Aves) categories. By comparing the mtDNA mutational spectrum (A_H_>G_H_ and A_H_>G_H_/ T_H_>C_H_) between these groups, we observed an expected trend: an increased A_H_>G_H_ and A_H_>G_H_ asymmetry in the warm-blooded classes (Fig. 4A, W = 430270, p-value = 5.878e-08), indicating that the mtDNA mutational spectrum is temperature sensitive in all vertebrates.

**FIG4.**
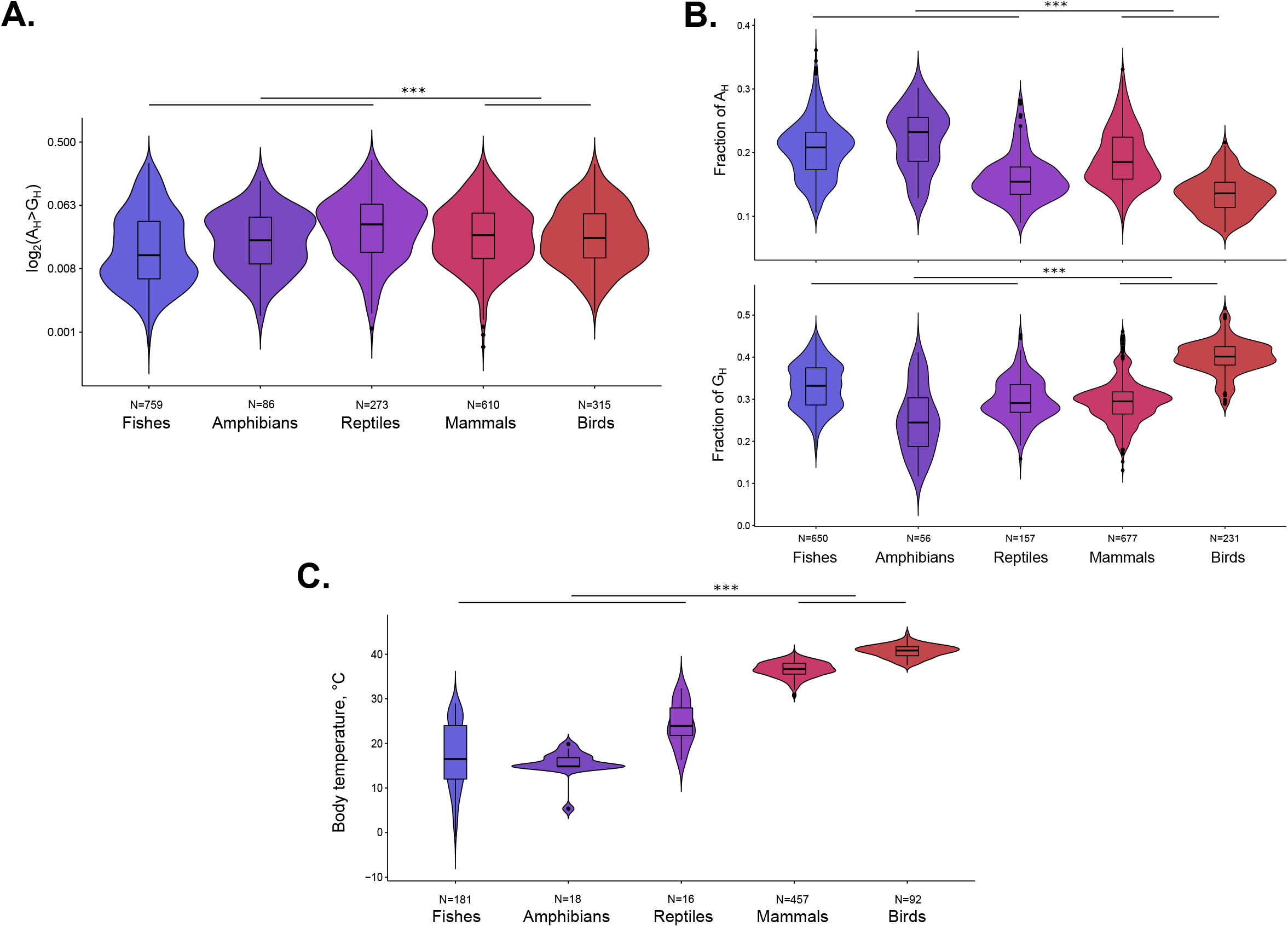
mtDNA mutational spectrum of all Vertebrates as a function of the average class-specific body temperature (the bottom panel). Between-class comparison shows an excess of A_H_>G_H_ (the first panel)and an excess of A_H_>G_H_/T_H_>C_H_ (the middle panel) in warm-versus cold-blooded species. All p-values < 2*10^-13^, Mann-Whitney U test.

Since the mtDNA mutational spectrum of all vertebrates goes in line with temperature (Fig. 4A), we may expect, that the whole genome neutral nucleotide content can also be shaped by the temperature-specific mutational bias: warm-blood species (mammals and birds) can be more A_H_-poor and G_H_-rich as compared to cold-blood ones. Using nucleotide content in fourfold degenerate sites of 12 genes (all but ND6), we indeed observed a deficit of A_H_ and an excess of G_H_ in warm-versus cold-blooded vertebrates (Fig. 4B, W = 0, all p-values < 2.2e-16). This result suggests the potentially broad effect of temperature (Fig. 4C, W = 14, p-value < 2.2e-16), which in the near future might be described for other taxa.

## DISCUSSION

By using an evolutionary approach, we were able to reconstruct a large number of mtDNA mutations in vertebrate species. This dataset allowed us to compare mutational spectra between species with different life-history and physiological characteristics. While previous research has shown that there is variation in the mtDNA mutational spectrum between species (Belle et al. 2005), the cause of this variation has not been explained. In this study, we hypothesized that the vast majority of fourfold degenerate synonymous substitutions are effectively neutral, and that differences in mutational spectra therefore reflect different mutagens rather than selection. To our knowledge, there is no evidence of selection acting on synonymous codons in the mtDNA of vertebrates. In fact, research has shown that mutational bias shapes the synonymous codon usage of human mtDNA (Ju et al. 2014) and the mtDNA of other mammals (Mikhailova et al. 2022).

Polymerase-induced mutations are expected to be symmetrical and enriched in A_H_>G_H_ and T_H_>C_H_ (Lee and Johnson 2006). However, the asymmetrical nature of mtDNA mutations observed in our study and other research (Arbeithuber et al. 2020; Mikhailova et al. 2022), (Yuan et al. 2020; Ju et al. 2014) and the positive relationship between A_H_>G_H_ and temperature and longevity requires an alternative explanation, potentially involving an additional mutagen that affects predominantly the single-stranded heavy chain during mtDNA replication (Fig S1.1). The temperature sensitivity of the mutational spectrum (Fig 1 - 4) may be due to increased oxidative damage in warm, highly-metabolic species (Martin and Palumbi 1993). To test the effect of oxidative damage more directly, we extracted experimental data on oxygen consumption (mg oxygen per kilogram fish per hour at 20°C at standard activity level) for a small number of fish species with known mutational spectra. Despite the extremely low sample size (N = 19), we observed a positive trend between oxygen consumption and A_H_>G_H_ (Supplementary Mat. 7). Therefore, the main finding of our paper - that temperature has an effect on the mutational spectrum - can be explained through the following logical steps: (i) temperature is linked to the level of aerobic metabolism, (ii) oxidative damage is a byproduct of aerobic metabolism, (iii) the asymmetry of AH>GH increases due to oxidative damage. Recent mutational accumulation experiments with *Chironomus riparius* and *E. coli* (Waldvogel and Pfenninger 2021; Chu et al. 2018) have shown that the fraction of transitions increases under high-temperature conditions. However, further experimental studies are needed to more fully understand the detailed mutational signatures of temperature.

The secondary result of our paper - longevity-associated changes in the mtDNA mutational spectrum is in line with our recent paper (Mikhailova et al. 2022). Longevity effect, i.e. relatively higher adenine deamination (increased A_H_>G_H_) in long-versus short-lived species, can be based on three non-mutually exclusive scenarios: (i) increased oxidative damage in oocytes of long-lived species, (ii) increased impact of mtDNA replication-mediated mutations in short-lived species and (iii) increased fidelity of mtDNA polymerase of long-lived species. The first scenario is supported by the recent experimental discovery that aged mouse oocytes are characterized by significantly increased A_H_>G_H_ as compared to young mouse oocytes (Arbeithuber et al.2020). An extension of the results from young and old mouse oocytes to short- and long-lived species will bring us to the observed correlations (Fig 4, 6) due to age-associated oxidative damage. The second scenario assumes that short-lived species per unit of time have an excess of replication-driven mtDNA mutations, which are expected to be symmetrical (A_H_>G_H_ ~ T_H_>C_H_, Supplementary Fig 1.1). This replication-driven input increases the overall symmetry of spectra diminishing the relative effect of asymmetrical oxidative damage-induced mutations in short-lived species. The third scenario is similar with the second one since it is associated with an excess of symmetrical replication-driven mutations in short-lived species, however it is based on higher fidelity of mtDNA polymerase in long-lived species (Nabholz, Glémin, and Galtier 2008), which could evolve to support decelerated somatic mutagenesis of mtDNA in long-lived large-bodied species.

Taking into account that all our observations (effects of temperature and longevity) might be explained through the same mediator - oxidative damage, which facilitates A>G substitutions on a single stranded heavy chain of mtDNA, we consider the oxidative damage explanation as the most parsimonious one. In other words, we propose that oxidative mutational signature in mtDNA is a function of both: metabolism-associated damage (approximated by temperature) and aging-associated damage (approximated by longevity).

The effect of temperature is pleiotropic - it affects not only the level of aerobic metabolism and oxidative damage but also life-history traits, such as generation time (Waldvogel and Pfenninger 2021): indeed high temperature is associated with short generation length in our (piece of results) and other studies. Thus the mutational signature of temperature is expected to be at least double. Some mutations are more sensitive to oxidative damage, while others are more replication-driven and depend rather on the rate of mtDNA turnover. Using our and other results, we propose that A_H_>G_H_ is mainly an oxidative damage-driven substitution, while C_H_>T_H_ is mainly a replication-driven one. As an overall working hypothesis, we propose, that temperature-related increase in oxidative damage is marked by the A_H_>G_H_, A_H_>G_H_ asymmetry, and G_H_A_H_ skew (a piece of results); while the temperature-related decreased generation length, which affects the mtDNA turnover rate, is marked by the C_H_>T_H_ and T_H_C_H_skew (TCskew depends on both temperature and maturation time: a piece of results).

Here there is an interesting analogy with gender-specific de novo mutational spectra in humans: maternally-inherited mutations in the nuclear genome resemble the oxidative damage (postmitotic female germ line), while paternally-inherited ones (proliferative male germ line) resemble the replication-driven mutations ().

Interestingly, an absence of the C>T correlation with generation length in mammals (Mikh) can be explained by the fact that mammalian oocytes are postmitotic and accumulate mainly the damage-driven mutations (A>G), while in fishes oocytes continue to divide the whole life, accumulating both: damage-driven mutations (A>G) and replication-driven ones (C>T).

Thermophiles. Selection or mutagenesis. The strongest effect (the highest slope is always on GC4). Criticism of selection ()

WHY LONGEVITY DOESN’T WORK (oocytes divide the whole life - Alya’s link). - discussion? oocyte are proliferative and mtDNA genome keeps under stronger selection - no pronounced affects of cellular aging. ?? temperature instead of longevity for A>G? Because there is no age-dependent damage - proliferating oocytes. Damage is more connected to temperature.

It is interesting to discuss - why A>G is associated with generation length in mammals, while it is associated with temperature in fishes and don’t show any association with the time of maturation. Probably, effect of temperature on oxidative damage is much stronger and thus, the maturation time effect is becoming noisy and nonsignificant for fishes, while for mammals with relatively stable temperature, the generation time effect is becoming visible. additional explanation can be based on the fact that aging of fish oocytes is not associated with strong oxidative damage because female germ line is proliferative and thus, accumulation of damage-induced mutations is less pronounced in fishes as compared to mammals. It is logical to assume that postmitotic cells can be more sensitive to oxidative damage - for example any somatic mutations or modification within the nuclear genome will destry respiratory chain and will affect the mtDNA mutations.

https://sciwheel.com/work/item/4248945/resources/10217061/pdf

aging-associated oxidative damage is expected to be more pronounced in postmitotic tissues as female germline in mammals.

An oxidative damage is a normal by-product of aerobic metabolism and the higher the metabolism, the higher the damage is expected. However, numerous antioxidant protective mechanisms may evolve in parallel with the increased level of the metabolism, partially or completely compensating the increasing oxidative damage, and thus decreasing the mutational rate. Observed in our paper correlations between several components of the mutational spectrum and temperature confirms the direct logic: the higher the metabolism, the higher the damage, demonstrating that the antioxidant protective mechanisms even if they compensate the oxidative damage, do it partially. This suggests that on the comparative-species scale, the fluctuations in the oxidative damage can be an effectively neutral trait not significantly affecting fitness of the analysed species.

A sensitivity of mtDNA mutational spectrum to environmental and life-history traits opens a possibility to use this metric in ecological and evolutionary studies. The reconstruction of the mtDNA mutational spectrum from neutral segregating polymorphisms of each species is rather straightforward and simple which can shed light on species-specific traits. For example, we ranked all analyzed Actinopterygii species by A_H_>G_H_ and compared species from the bottom and top deciles. Expectedly we observed that top decile (with high A_H_>G_H_) consists of many large-bodied freshwater species inhabiting predominantly tropic lakes and rivers: *Labeo gonius,* which spend a lot of time in Tropical climate zone; *Brachyhypopomus occidentalis* frequently found in shallow and calm streams; *Petrochromis trewavasae,* who lives in african *Lake Tanganyika; Girardinus metallicus* who lives in ponds, lakes and streams of Cuba in Central America; reef fish, such as *Mugil curema, Amblyeleotris wheeleri* and *Beaufortia kweichowensis,* living in shallow waters with oxygen-saturated water; *Lepisosteus oculatus* which is capable to compensate for the lack of oxygen by gulping air bubbles into a primitive lung called a gas bladder. Contrary, species from the bottom decile (with low A_H_>G_H_) are mainly small-sized sea or deepwater lake fishes such as: *Oncorhynchus mykiss,* which inhabit clear, cold headwaters, creeks, small to large rivers, lakes, and intertidal areas, usually not stocked in water reaching summer temperatures above 25°C or ponds with very low oxygen concentrations; *Hippoglossoides robustus,* that lives on bottoms at depths of up to 425 metres where temperature reach up to 8°C, *Comephorus dybowskii* from Baikal lake found beyond 1,000 m depth.

Indeed, it has been shown experimentally on mouse cells (Shin and Turker 2002) and bacteria (Shewaramani et al. 2017) that oxidative damage predominantly increases adenine deamination especially on single-stranded DNA. Thus, higher adenine deamination (A_H_>G_H_) in species with increased temperature can be explained by their higher oxidative damage due to intensive aerobic metabolism.

## Supporting information

Fupplementary File 1

## SUPPLEMENTARY FILES

Supplementary file 1 A complete collection of 12-component mtDNA mutational spectra for 551 Actinopterygii species, used in the current study. For Actinopterygii species added known ambient temperature AND time of maturation

## METHODS

The widely accepted annotation of mitochondrial genomes is based on the light chain, however it is known that the majority of mitochondrial mutations occur on a heavy chain (Faith and Pollock 2003). In order to emphasize the chemical nature of observed mutations, we presented all substitutions with heavy-chain notation.

The mutational spectrum reconstruction has been described in detail in our accompanying paper (Mikhailova et al. 2022). Here, we describe the logic of the analysis briefly. The mutational spectrum (a probability of one nucleotide to mutate into any other nucleotide) for each Vertebrate species was derived from all available intraspecies sequences (as of April 2018) of mitochondrial protein-coding genes. Using this database with intraspecies polymorphisms and developed pipeline we reconstructed the intraspecies phylogeny using an outgroup sequence (closest species for analyzed one), reconstructed ancestral states spectra in all positions at all inner tree nodes and finally got the list of single-nucleotide substitutions for each gene of each species. Using species with at least 15 single-nucleotide synonymous mutations at four-fold degenerate sites we estimated the mutational spectrum for more than a thousand chordata species. We normalized observed frequencies by nucleotide content in the third position of four-fold degenerative synonymous sites of a given gene. Details of the pipeline are described in our recent paper (Mikhailova et al. 2022)and the separate methodological paper is under preparation.

All statistical analyses were performed in R using Spearman rank correlations and Multiple models (for mammalian analyses dummy variables for each group were used). PGLS method (package “caper”, version 1.0.1) was used for the analysis of phylogenetic inertia.

The annual mean environmental (water) temperature in Celsius and time of maturation in years (mean or median age at first maturity, at which 50% of a cohort spawn for the first time) for fishes were downloaded from https://www.fishbase.se/ (at September 2019).

Mutational spectra of the European anchovy clades were estimated using a simplified algorithm, as compared to the between-species analyses described above. First, for each clade (285 organisms of clade A and 160 organisms of clade B, provided by Dr. Rita Castilho) we reconstructed a consensus of the region of MT-CYB, assuming that consensus reflects the ancestral state. Second, within each clase we counted all synonymous deviations from consensus (irrespectively of the frequency of the minor allele - from singletons to alleles with MAF < 50%), splitting these SNPs into five groups: four types of transitions and a group of all transversions (as in Fig 2) which were used in the analyses. We didn’t normalize the observed substitutions by the nucleotide content taking into account that compared sequences and clades are very similar.

S_TG_-S_AC_ was calculated as a difference between sums of pairs of relative nucleotide frequencies: (sum(FrT+FrG) - sum(FrA+FrC))/(sum(FrA+FrC) + sum(FrT+FrG)), where (sum(FrA+FrC) + sum(FrT+FrG) = 1.

## ACKNOWLEDGMENTS

D. I. was supported by the federal academic leadership program Priority 2030 at the Immanuel Kant Baltic Federal University. E.O.T. is supported by a scholarship from the Austrian Science Fund (FWF, DOC 33-B27). A.G.M., K.P. and V.Y. were supported by the Russian Science Foundation grant No. 21-75-20143. I.M. was supported by the Russian Science Foundation No. 21-75-10081. K.G. was supported by the Russian Science Foundation grant No. 21-75-20145. A. K. was supported by the Ministry of Science and Higher Education of the Russian Federation (agreement no. 075-15-2021-1084).

