## Supplementary Materials for "A>G substitutions on a heavy chain of mitochondrial genome marks an increased level of aerobic metabolism in warm versus cold vertebrates"

Supplementary Materials to the manuscript by Mikhailova et al. “A>G substitutions on a heavy chain of mitochondrial genome marks an increased level of aerobic metabolism in warm versus cold vertebrates .”

### 1. Species-specific mtDNA mutational spectrum of fishes is associated with ambient water temperature by means of increased $A_H>G_H$ and $A_H>G_H$ asymmetry.

***https://github.com/mitoclub/MutSpecOfActinopterygii/blob/master/Scripts/1.VertebratePolymorphicData.Actinopterygii.Analyses.html***

**a. Spearman rank correlations of the two most common transitions and their asymmetries with temperature. Frequencies of all other types of substitutions didn’t show significant correlations with temperature (p-values > 0.1):**

| ***taxa*** | ***substitution*** | ***Spearman’s Rho*** | ***p value*** |
|---|---|---|---|
| fish<br>*N = 188* | $A_H>G_H$ | 0.2985488 | 3.162e-05 |
| | $A_H>G_H/T_H>C_H$ | 0.3889124 | 2.654e-07 |
| | $C_H>T_H$ | 0.1939811 | 0.007644 |
| | $C_H>T_H/G_H>A_H$ | -0.08912368 | 0.2564 |

**b. A linear model describing $A_H>G_H$ asymmetry as a function of temperature:**

| ***model*** | ***variable*** | ***coefficient*** | ***p value*** |
|---|---|---|---|
| $A_H>G_H$ ~ scale(Temperature)<br>*N = 188* | Intercept | 0.018959 | < 2e-16 *** |
| | scale(Temperature) | 0.007750 | 9.66e-05 *** |
| $A_H>G_H/T_H>C_H$ ~ scale(Temperature)<br>*N = 188* | Intercept | 3.6584 | <2e-16 *** |
| | scale(Temperature) | 0.9647 | 0.0145 * |
| $C_H>T_H$ ~ scale(Temperature)<br>*N = 188* | Intercept | 0.068390 | < 2e-16 *** |
| | scale(Temperature) | 0.015680 | 0.00638 ** |
| $C_H>T_H/G_H>A_H$ ~ scale(Temperature)<br>*N = 188* | Intercept | 10.1405 | <2e-16 *** |
| | scale(Temperature) | -1.0889 | 0.181 |

**c. Phylogenetic inertia analyses (phylogenetic generalised least-squares (PGLS) regression model) of $A_H>G_H$ asymmetry as a function of temperature**

| ***model*** | ***variable*** | ***coefficient*** | ***p value*** |
|---|---|---|---|
| log2($A_H>G_H$) ~ log2(Temperature)<br>*N = 52, lambda [ ML] : 0.000* | Intercept | -8.91589 | < 2.2e-16 *** |
| | log2(Temperature) | 0.70605 | 0.0002791 *** |
| log2($A_H>G_H/T_H>C_H$) ~ log2(Temperature)<br>*N = 52, lambda [ ML] : 0.000* | Intercept | -0.59919 | 0.341474 |
| | log2(Temperature) | 0.44507 | 0.009124 ** |
| log2($C_H>T_H$) ~ log2(Temperature)<br>*N = 52, lambda [ ML] : 0.667* | Intercept | -5.39515 | 6.384e-07 *** |
| | log2(Temperature) | 0.39050 | 0.02828 * |

| | | | |
|---|---|---|---|
| $\log_2(C_H{>}T_H/G_H{>}A_H)$ ~ log2(Temperature)<br>*N = 52, lambda [ ML] : 0.000* | Intercept | 3.25550 | 1.982e-07 *** |
| | log2(Temperature) | -0.12593 | 0.3795 |

**Supplementary Figure 1.1 Asymmetry of equivalent substitutions traces a mutagen (potentially an oxidative damage) predominantly affecting a heavy chain.**

If a mutagenic process introduces for example mutation A>G on one chain it will be equivalent to T>C on the opposite chain and thus symmetrical mutagenesis would lead to similar probabilities of A>G and T>C on any chain. However, probabilities are not equal with $A_H{>}G_H$ being significantly higher than $T_H{>}C_H$ (Fig 1A). This means that some mutagen(s) affects predominantly heavy chain, which stays single-stranded during mtDNA replication, introducing $A_H{>}G_H$ substitutions. An increased temperature is associated with even more elevated asymmetry: $A_H{>}G_H$ /$T_H{>}C_H$ (Fig 1B). We interpret the increased asymmetry of $A_H{>}G_H$ /$T_H{>}C_H$ in warm-water species as an increased damage of the heavy chain of mtDNA due to high temperature (high level of aerobic metabolism and, correspondingly higher oxidative damage).

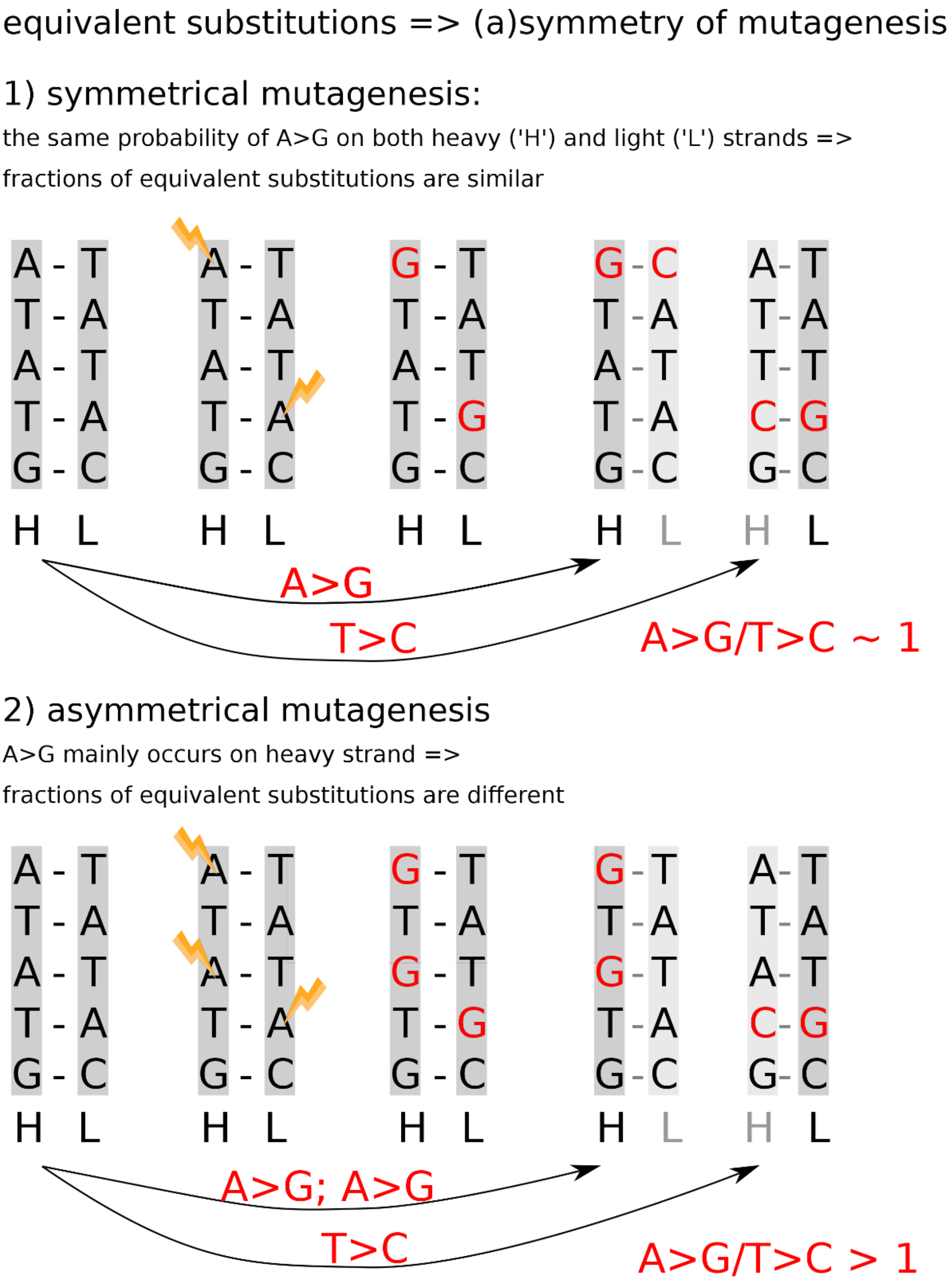

**Supplementary Figure 1.2 mtDNA spectrum of fishes is associated with temperature: family--specific analyses**

***https://github.com/mitoclub/MutSpecOfActinopterygii/blob/master/Scripts/AnalysGraphsForSupplements.Rmd***

Median family-specific $A_H>G_H$ asymmetry ($A_H>G_H/T_H>C_H$) is associated positively with median family-specific temperature (N = 19, p = 0.02631 Rho = 0.51)

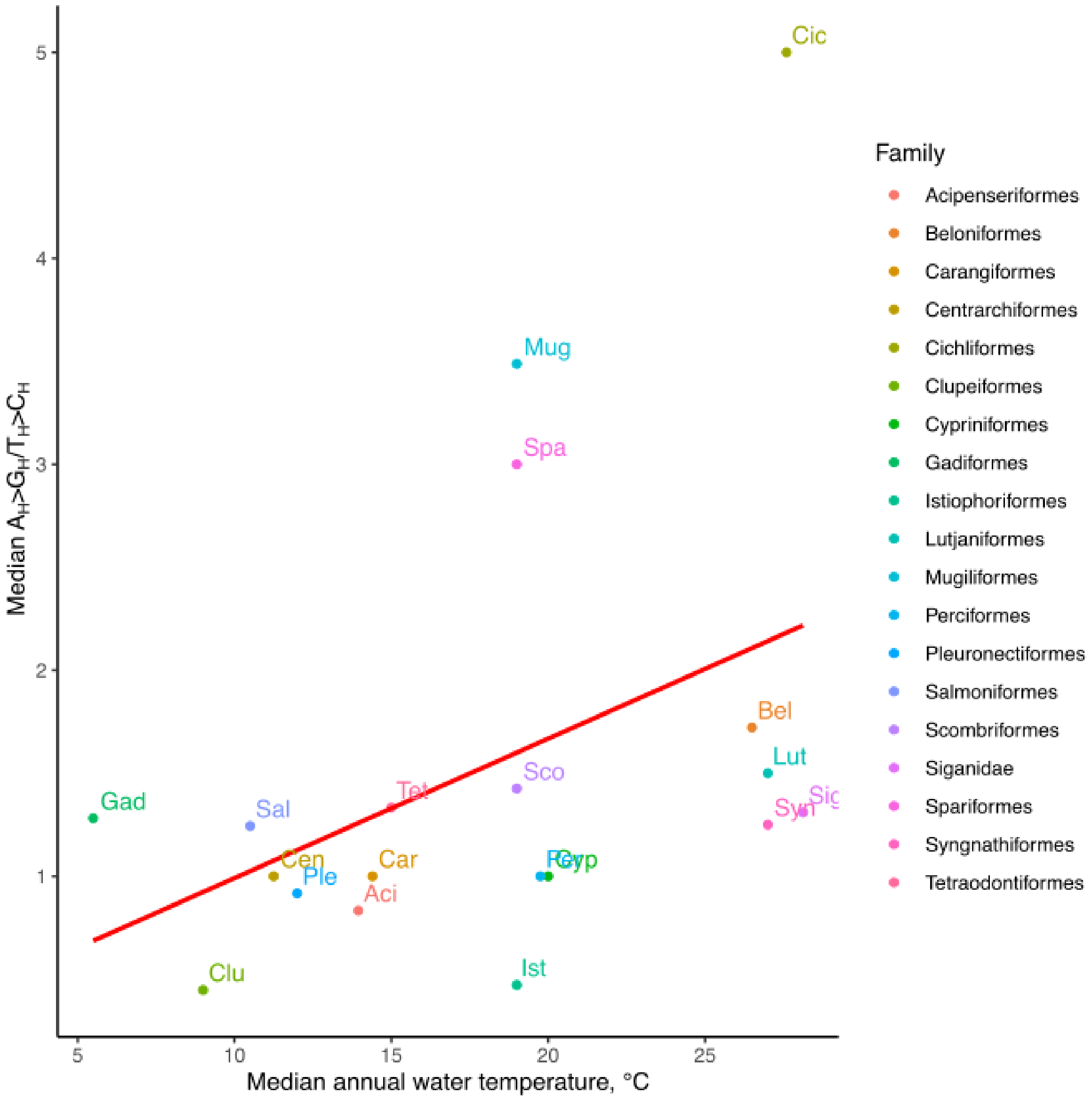


**d. Spearman rank correlation between temperature and the time of maturation in fishes**

| ***taxa*** | ***Spearman's Rho*** | ***p value*** |
|---|---|---|
| fishes<br>*N = 96* | -0.3287072 | 0.001076 |

**e. Spearman rank correlations between the time of maturation and the key components of the mutational spectrum: $A_H>G_H$, $C_H>T_H$, $A_H>G_H$ asymmetry ($A_H>G_H/T_H>C_H$) and $C_H>T_H$ asymmetry ($C_H>T_H/G_H>A_H$).**

| *taxa* | *variable* | *Spearman's Rho* | *p value* |
|---|---|---|---|
| fishes<br>*N = 96* | $A_H>G_H$ | -0.1569224 | 0.1268 |
| fishes<br>*N = 96* | $C_H>T_H$ | 0.005575674 | 0.957 |
| fishes<br>*N = 96* | $A_H>G_H/T_H>C_H$ | -0.04573348 | 0.6229 |
| fishes<br>*N = 96* | $C_H>T_H/G_H>A_H$ | -0.0616844 | 0.5051 |

**f. Multiple linear models with $A_H>G_H$ and $C_H>T_H$ and $A_H>G_H$ asymmetry ($A_H>G_H/T_H>C_H$) and $C_H>T_H$ asymmetry ($C_H>T_H/G_H>A_H$) as functions of the temperature and the time of maturation:**

| *model* | *variable* | *coefficient* | *p value* |
|---|---|---|---|
| A) $A_H>G_H$ ~ scale(Temperature) * scale(TimeOfMaturation)<br>*N= 96* | Intercept | 0.020323 | 2.45e-09 *** |
| | scale(Temperature) | 0.009526 | 0.0024 ** |
| | scale(TimeOfMaturation) | -0.001981 | 0.5623 |
| | scale(Temperature): scale(TimeOfMaturation) | -0.001094 | 0.7280 |
| B) $A_H>G_H$~ scale(Temperature) + scale(TimeOfMaturation)<br>*N= 96* | Intercept | 0.020647 | 2.68e-10 *** |
| | scale(Temperature) | 0.009511 | 0.00232 ** |
| | scale(TimeOfMaturation) | -0.001431 | 0.63500 |
| A) $A_H>G_H/T_H>C_H$ ~ scale(Temperature) * scale(TimeOfMaturation)<br>*N= 69* | Intercept | 3.6506 | 5.35e-12 *** |
| | scale(Temperature) | 1.0256 | 0.0168 * |
| | scale(TimeOfMaturation) | 0.5740 | 0.2578 |
| | scale(Temperature): scale(TimeOfMaturation) | 0.7080 | 0.1184 |
| B) $A_H>G_H/T_H>C_H$ ~ scale(Temperature) + scale(TimeOfMaturation)<br>*N= 69* | Intercept | 3.4250 | 8.86e-12 *** |
| | scale(Temperature) | 0.9908 | 0.0219 * |
| | scale(TimeOfMaturation) | 0.1439 | 0.7377 |
| A) $C_H>T_H$ ~ scale(Temperature) * scale(TimeOfMaturation)<br>*N= 96* | Intercept | 0.074275 | 1.47e-11 *** |
| | scale(Temperature) | 0.013601 | 0.159 |
| | scale(TimeOfMaturation) | 0.004570 | 0.670 |

| | | | |
|---|---|---|---|
| | scale(Temperature): scale(TimeOfMaturation) | 0.006202 | 0.530 |
| B) $C_H$>$T_H$ ~ scale(Temperature) + scale(TimeOfMaturation) *N= 96* | Intercept | 0.072437 | 5.25e-12 *** |
| | scale(Temperature) | 0.013687 | 0.155 |
| | scale(TimeOfMaturation) | 0.001455 | 0.878 |
| A) $C_H$>$T_H$/$G_H$>$A_H$ ~ scale(Temperature) * scale(TimeOfMaturation) *N= 69* | Intercept | 9.5799 | 1.36e-10 *** |
| | scale(Temperature) | 0.2455 | 0.840 |
| | scale(TimeOfMaturation) | 1.0112 | 0.490 |
| | scale(Temperature): scale(TimeOfMaturation) | 0.5267 | 0.686 |
| B) $C_H$>$T_H$/$G_H$>$A_H$ ~ scale(Temperature) + scale(TimeOfMaturation) *N= 69* | Intercept | 9.4122 | 2.87e-11 *** |
| | scale(Temperature) | 0.2197 | 0.855 |
| | scale(TimeOfMaturation) | 0.6912 | 0.572 |

**g. Multiple linear models with temperature and the time of maturation as functions of $A_H$>$G_H$ and $C_H$>$T_H$**

| ***model*** | ***variable*** | ***coefficient*** | ***p value*** |
|---|---|---|---|
| Temperature ~ scale($A_H$>$G_H$) + scale($C_H$>$T_H$) *N= 188* | Intercept | 17.12660 | < 2e-16 *** |
| | scale($A_H$>$G_H$) | 2.05598 | 0.00544 ** |
| | scale($C_H$>$T_H$) | 0.05291 | 0.94237 |
| TimeOfMaturation ~ scale($A_H$>$G_H$) + scale($C_H$>$T_H$) *N= 188* | Intercept | 3.9298 | <2e-16 *** |
| | scale($A_H$>$G_H$) | -0.6647 | 0.101 |
| | scale($C_H$>$T_H$) | 0.3372 | 0.382 |

### 2. Tropical clade of European anchovy shows an excess of $A_H$>$G_H$ and deficit of $T_H$>$C_H$ in mtDNA

*https://github.com/polarsong/mtDNA_mutspectrum/blob/MutSpecOfActinopterygii/Head/2Scripts/2b.AnchoviesMutSpecAnalyses.FiguresStatistics.R*

**a. Differences between consensuses of clades A and B:**

*https://github.com/polarsong/mtDNA_mutspectrum/blob/MutSpecOfActinopterygii/Body/2Derived/Anchovies_Cytb_diff.csv*

| Position | Codon of clade A | Codon of clade B | Aminoacid of clade A | Aminoacid of clade B |
|---|---|---|---|---|
| 131 | GGG | GGA | G | G |
| 137 | CTT | CTA | L | L |
| 173 | TTT | TTC | F | F |
| 185 | ATT | ATC | I | I |
| 203 | CGG | CGA | R | R |
| 383 | TTG | TTA | L | L |
| 386 | GTC | GTG | V | V |
| 389 | CAA | CAG | Q | Q |
| 455 | TTG | TTA | L | L |
| 467 | GTC | GTT | V | V |
| 530 | GCG | GCA | A | A |
| 596 | GGG | GGA | G | G |
| 617 | GCA | GCG | A | A |
| 626 | TCA | TCG | S | S |
| 638 | TTT | TTC | F | F |
| 719 | TGG | TGA | W | W |
| 749 | CGG | CGA | R | R |
| 752 | TCT | TCC | S | S |
| 758 | CCG | CCA | P | P |
| 770 | GGT | GGC | G | G |
| 797 | ATT | ATC | I | I |

### 3. The temperature sensitive mtDNA mutational spectrum of ray-finned fishes affects their neutral nucleotide composition: warm water species are more AC-poor and GT-rich

#### a. Typical mitochondrial mutational spectra of cold- and warm- water fishes.

| | $T_H$_$A_H$ | $T_H$_$C_H$ | $T_H$_$G_H$ | $A_H$_$T_H$ | $A_H$_$C_H$ | $A_H$_$G_H$ | $C_H$_$T_H$ | $C_H$_$A_H$ | $C_H$_$G_H$ | $G_H$_$T_H$ | $G_H$_$A_H$ | $G_H$_$C_H$ |
|---|---|---|---|---|---|---|---|---|---|---|---|---|
| **cold fish** | 0.009173495 | 0.08326 | 0.006509080 | 0.012862530 | 0.003601526 | 0.100910211 | 0.634310246 | 0.009593790 | 0.023799040 | 0.031747375 | 0.078514468 | 0.006617838 |
| **warm fish** | 0.004864868 | 0.099896169 | 0.011772136 | 0.023065033 | 0.009870074 | 0.179538135 | 0.459716445 | 0.014697295 | 0.022697971 | 0.024094913 | 0.137717075 | 0.012069886 |

**Supplementary Figure 3.1 Expected nucleotide content at compositional equilibrium:**

***https://github.com/mitoclub/MutSpecOfActinopterygii/blob/master/Scripts/EqulibriumFishes.Rmd***

The typical mutational spectra for cold- and warm- water fishes was derived based on a subset of the coldest and warmest species (the coldest decile, N = 19; the warmest decile, N = 23). Next, using these spectra we derived an expected nucleotide content at compositional equilibrium. In order to do it we used two approaches: simulations (R script) and differential equations (equations are available upon request and the method paper is under preparation). Briefly, the main goal of simulations was to obtain an equilibrium nucleotide composition (frequencies of four nucleotides) evolved under a given mutational spectrum. Simulations started with random frequencies of four nucleotides (left part of the plots) and with time (with the simulated generations) the nucleotide frequencies changed according to the mutational spectrum, reaching the saturation (right part of the plots). Both approaches demonstrated identical expected nucleotide composition (see below: solid curves correspond to the simulations which converge during the simulation at one value, dotted lines correspond to the analytical solution, obtained as a solution of differential equations ). The equilibrium values (4 nucleotide frequencies for each temperature) were used in our downstream analyses as expected nucleotide content (see Fig 3A in the main text for details).

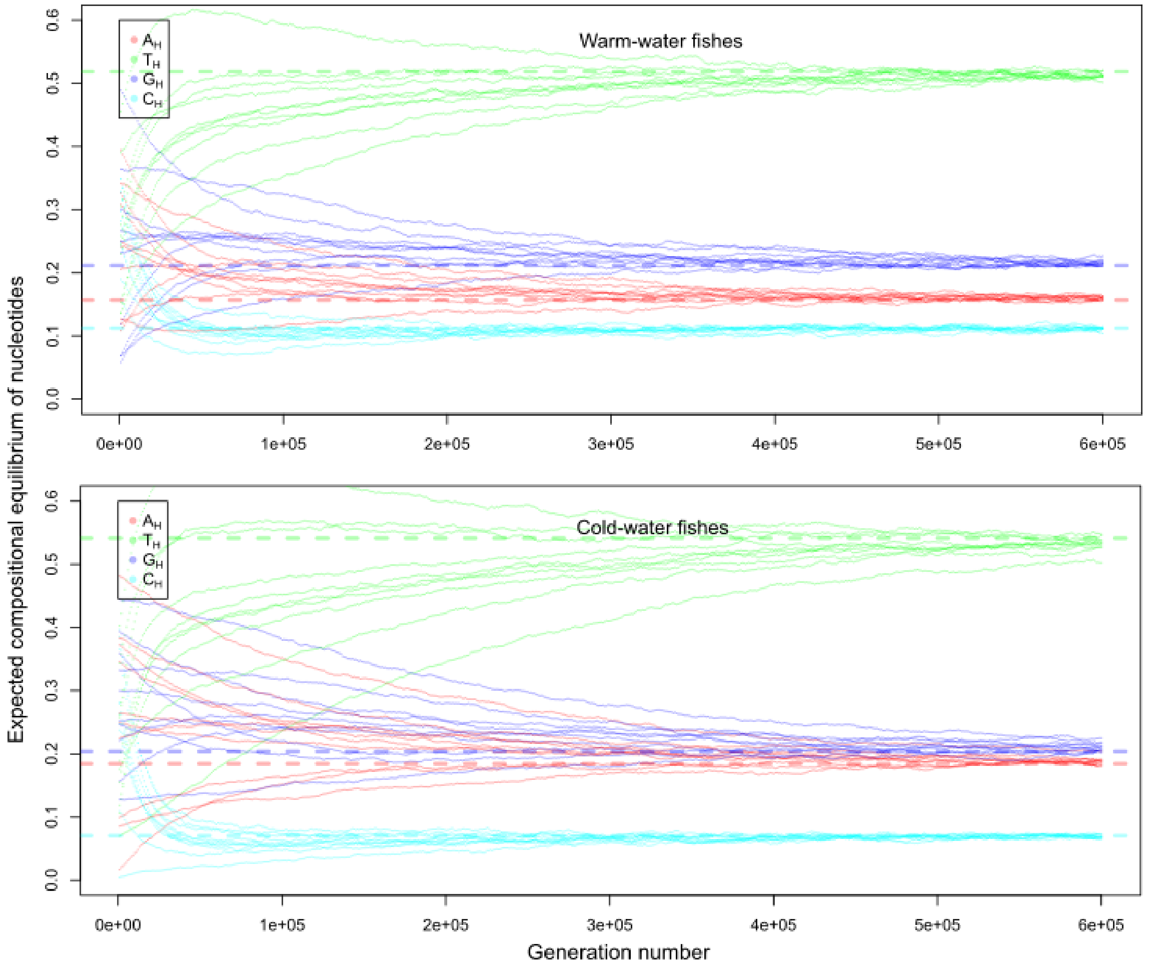

***https://github.com/mitoclub/MutSpecOfActinopterygii/blob/master/Scripts/3.WholeGenomeDataAnalyses.Actinopterygii.Analyses.html***

**b. Spearman rank correlations between fractions of 4 nucleotides, GAskew, TCskew and temperature:**

| *class* | *Fraction of nucleotide* | *Spearman's Rho* | *p-value* |
|---|---|---|---|
| Actinopterygii *N = 333* | $A_H$ | -0.1612848 | 0.003163 |
| | $T_H$ | 0.1318643 | 0.01605 |
| | $G_H$ | 0.1235587 | 0.02414 |
| | $C_H$ | -0.2548114 | 2.473e-06 |
| | GAskew | 0.1816663 | 0.0008673 |
| | TCskew | 0.2429729 | 7.307e-06 |

**c. Spearman rank correlations between fractions of 4 nucleotides, GAskew, TCskew and time of maturation:**

| *class* | *Fraction of nucleotide* | *Spearman's Rho* | *p-value* |
|---|---|---|---|
| Actinopterygii *N = 217* | $A_H$ | -0.06058898 | 0.3744 |
| | $T_H$ | 0.2597488 | 0.0001084 |
| | $G_H$ | -0.06446041 | 0.3446 |
| | $C_H$ | -0.1890223 | 0.005212 |
| | GAskew | 0.003450675 | 0.9597 |
| | TCskew | 0.2322227 | 0.000564 |

**d. Linear regression models between GAskew, TCskew, $S_{TG}$-$S_{AC}$ and temperature and time of maturation:**

| *model* | *variable* | *coefficients* | *p values* |
|---|---|---|---|
| GAskew ~ Temperature *N=333* | Intercept | 0.174308 | 8.01e-14 *** |
| | Temperature | 0.002867 | 0.00802 ** |
| TCskew ~ Temperature *N=333* | Intercept | 0.6054590 | < 2e-16 *** |
| | Temperature | 0.0036119 | 2.63e-05 *** |
| $S_{TG}$-$S_{AC}$ ~ Temperature *N=333* | Intercept | 0.3576532 | < 2e-16 *** |
| | Temperature | 0.0034870 | 3.22e-07 *** |
| GAskew ~ TimeOfMaturation *N=217* | Intercept | 0.1876786 | 7.27e-16 *** |
| | TimeOfMaturation | 0.0007169 | 0.821 |
| TCskew ~ TimeOfMaturation *N=217* | Intercept | 0.659894 | < 2e-16 *** |
| | TimeOfMaturation | 0.006972 | 0.00534 ** |
| $S_{TG}$-$S_{AC}$ ~ TimeOfMaturation *N=217* | Intercept | 0.390449 | <2e-16 *** |

| | | | |
|---|---|---|---|
| | TimeOfMaturation | 0.004200 | 0.0234 * |

**e. Phylogenetic inertia analyses (phylogenetic generalised least-squares (PGLS) regression model):**

| ***model*** | ***variable*** | ***coefficients*** | ***p values*** |
|---|---|---|---|
| GAskew ~ Temperature<br>*N=131, lambda [ ML] : 1.000* | Intercept | 0.1158136 | 0.2910 |
| | Temperature | 0.0015013 | 0.1548 |
| TCskew ~ Temperature<br>*N=131, lambda [ ML] : 0.979* | Intercept | 0.76067668 | <2e-16 *** |
| | Temperature | 0.00073197 | 0.5089 |
| $S_{TG}$-$S_{AC}$ ~ Temperature<br>*N=131, lambda [ ML] : 0.991* | Intercept | 0.40049676 | 1.629e-09 *** |
| | Temperature | 0.00130582 | 0.09781 . |

**f. Linear regression models between GAskew, TCskew, $S_{TG}$-$S_{AC}$ and temperature + the time of maturation:**

| | | | |
|---|---|---|---|
| GAskew ~ scale(Temperature + 2) + scale(TimeOfMaturation)<br>*N = 131* | Intercept | 0.20311 | < 2e-16 *** |
| | log2(Temperature + 2) | 0.04612 | 0.00233 ** |
| | log2(TimeOfMaturation) | 0.01452 | 0.32360 |
| TCskew ~ scale(Temperature + 2) + scale(TimeOfMaturation)<br>*N = 131* | Intercept | 0.70170 | < 2e-16 *** |
| | scale(Temperature + 2) | 0.04000 | 0.000641 *** |
| | scale(TimeOfMaturation) | 0.04201 | 0.000292 *** |
| $S_{TG}$-$S_{AC}$ ~ scale(Temperature + 2) + scale(TimeOfMaturation)<br>*N = 131* | Intercept | 0.421937 | < 2e-16 *** |
| | scale(Temperature + 2) | 0.047876 | 1.10e-08 *** |
| | scale(TimeOfMaturation) | 0.031083 | 9.82e-05 *** |

**g. Phylogenetic inertia analyses (phylogenetic generalised least-squares (PGLS) regression model)**

| ***model*** | ***variable*** | ***coefficients*** | ***p values*** |
|---|---|---|---|
| GAskew ~ log2(Temperature + 2) + log2(TimeOfMaturation)<br>*N=131, lambda [ ML] : 1.000* | Intercept | 0.0256617 | 0.83270 |
| | log2(Temperature + 2) | 0.0201862 | 0.08424 . |
| | log2(TimeOfMaturation) | 0.0115187 | 0.10653 |
| TCskew ~ log2(Temperature + 2) + log2(TimeOfMaturation)<br>*N=131, lambda [ ML] : 0.975* | Intercept | 0.7126050 | 4.671e-12 *** |
| | log2(Temperature + 2) | 0.0120913 | 0.3161 |
| | log2(TimeOfMaturation) | 0.0038918 | 0.5652 |
| $S_{TG}$-$S_{AC}$ ~ log2(Temperature + 2) + log2(TimeOfMaturation)<br>*N=131, lambda [ ML] : 0.986* | Intercept | 0.3198276 | 1.491e-05 *** |
| | log2(Temperature + 2) | 0.0184000 | 0.03428 * |
| | log2(TimeOfMaturation) | 0.0097012 | 0.04748 * |

**h. Linear regression models between temperature and GAskew + TCskew, the time of maturation and GAskew + TCskew:**

| | | | |
|---|---|---|---|
| Temperature ~ scale(GAskew) + scale(TCskew) *N = 131* | Intercept | 19.2908 | < 2e-16 *** |
| | scale(GAskew) | 1.5234 | 0.000278 *** |
| | scale(TCskew) | 2.0620 | 1.06e-06 *** |
| TimeOfMaturation ~ scale(GAskew) + scale(TCskew) *N = 131* | Intercept | 4.4415 | < 2e-16 *** |
| | scale(GAskew) | 0.4793 | 0.21477 |
| | scale(TCskew) | 1.1892 | 0.00244 ** |

**i. Phylogenetic inertia analyses (phylogenetic generalised least-squares (PGLS) regression model)**

| ***model*** | ***variable*** | ***coefficients*** | ***p values*** |
|---|---|---|---|
| Temperature ~ scale(GAskew) + scale(TCskew) *N=131, lambda [ ML] : 0.769* | Intercept | 14.73620 | 2.831e-05 *** |
| | scale(GAskew) | 2.10675 | 0.004665 ** |
| | scale(TCskew) | 2.46107 | 0.003183 ** |
| TimeOfMaturation ~ scale(GAskew) + scale(TCskew) *N=131, lambda [ ML] : 0.493* | Intercept | 8.754531 | 1.02e-06 *** |
| | scale(GAskew) | 1.078439 | 0.02547 * |
| | scale(TCskew) | 0.070535 | 0.89899 |

**Supplementary Figure 3.2 mtDNA nucleotide composition depends on temperature within both short- and long-maturated groups of Actinopterygii species**

*https://github.com/polarsong/mtDNA_mutspectrum/blob/MutSpecOfActinopterygii/Head/2Scripts/accessorialSCRIPTAnalysesGraphsForSupplements.R*

First, we split all fishes into 2 groups: short- and long-maturated, depending on the median of the time of maturation (median = 3 years). Second, each group was additionally divided into two subgroups: cold- and warm-water species based on a group-specific medians of temperature (19.7ºC and 15ºC for short and long-maturated groups correspondingly). We observed, that $S_{TG}$-$S_{AC}$ is higher in warm- versus cold-water fishes in both short maturated (p-value = 0.0002245, Mann-Whitney U test, 29 cold- versus 30 warm-water species) and long-maturated groups (p-value = 0.00105, Mann-Whitney U test, 35 cold- versus 37 warm-water species).

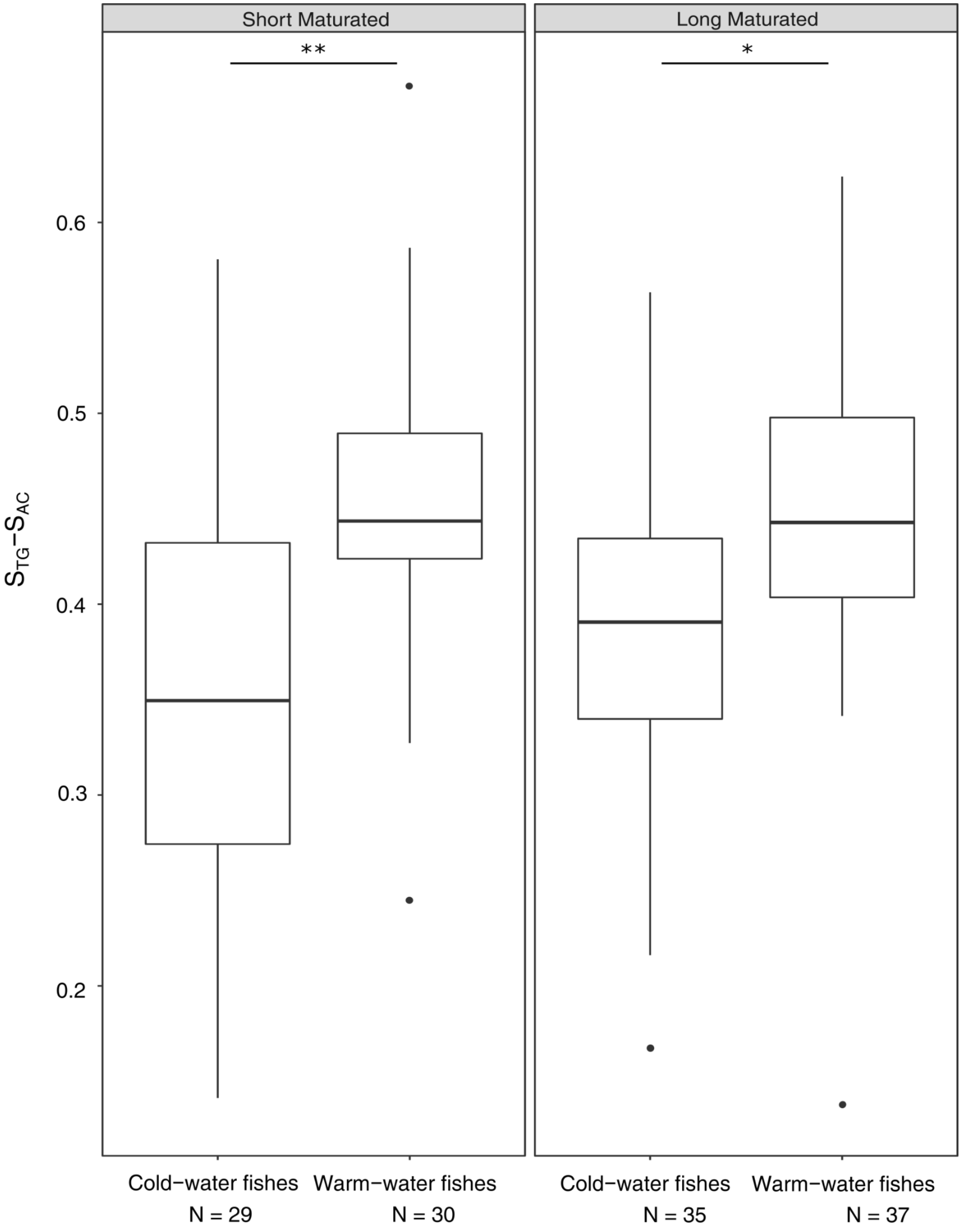

### 4. The mtDNA mutational spectrum is temperature sensitive in all classes of vertebrates

***https://github.com/mitoclub/MutSpecOfActinopterygii/blob/master/Scripts/4.VertebratePolymorphicData.ViolinPlot.Rmd***

Temperature was retrieved from AnAge database(https://genomics.senescence.info/species/index.html). Within each class subsets of species with known temperature and species with known mutational spectrum can differ from each other. However, due to the robust class-specific (not species-specific) comparisons we ignored this problem. Temperature of birds was retrieved from (https://doi.org/10.2307/1365174).

**Supplementary Figure 4**

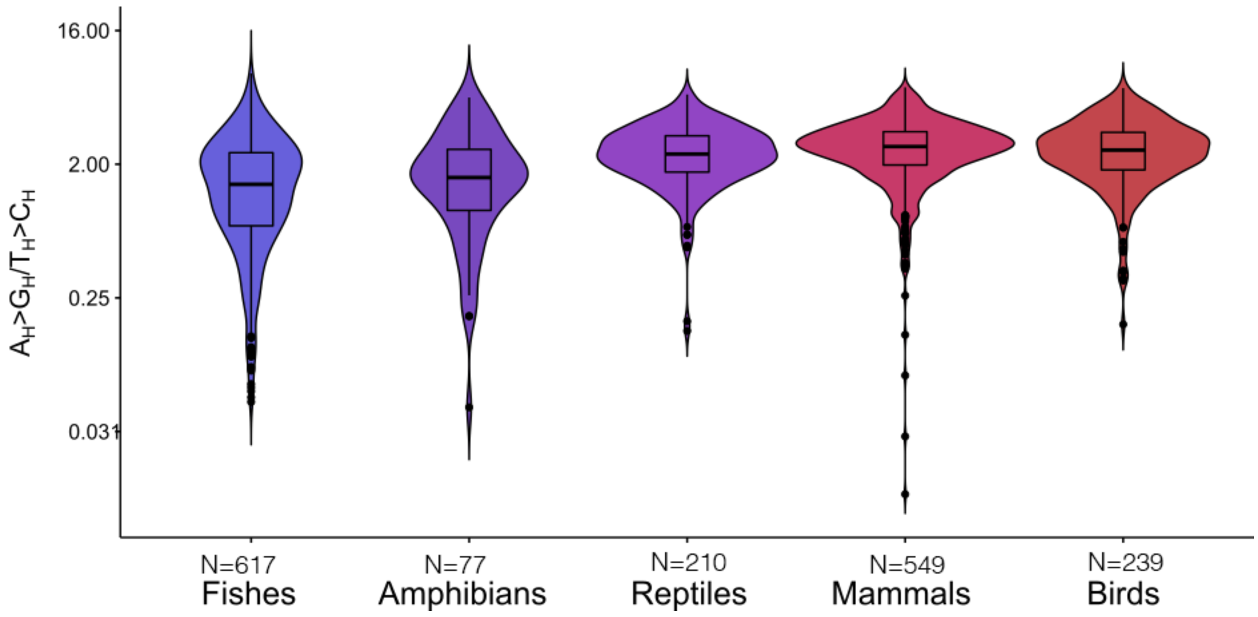


### 5. Oxygen consumption.

**Supplementary Figure 5 Oxygen consumption.**

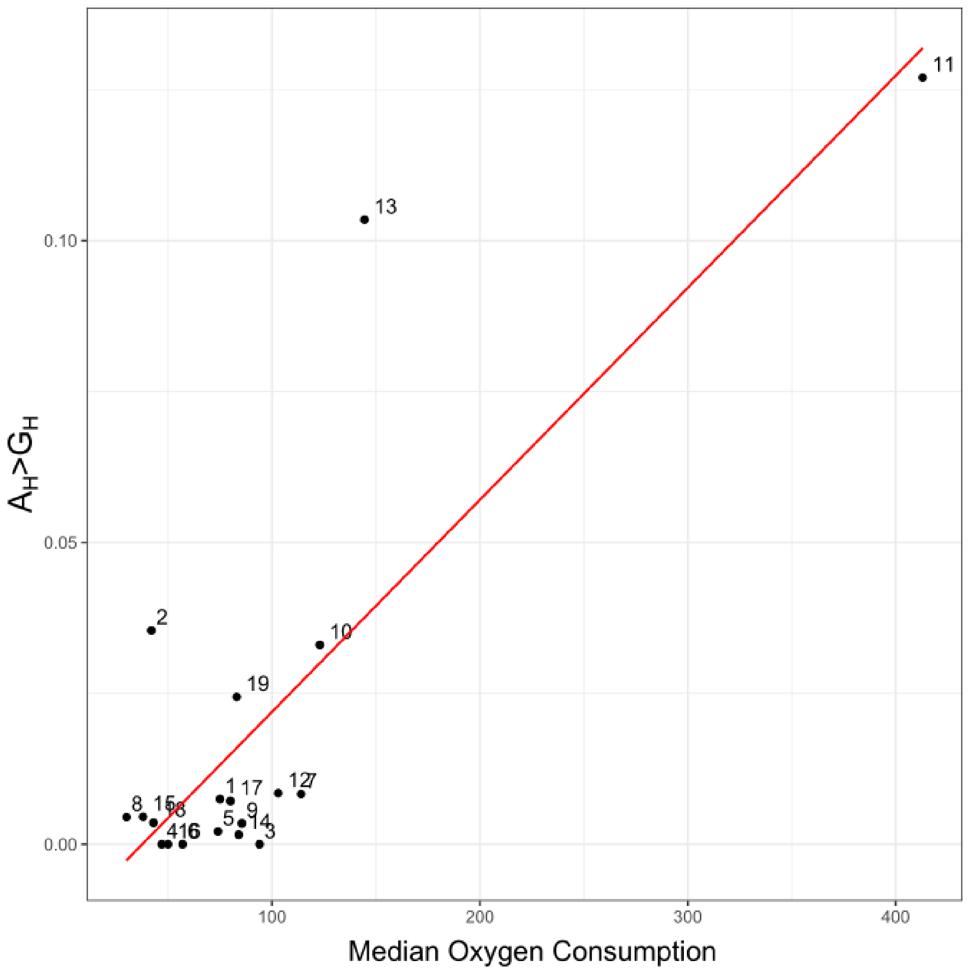

| Correlation Test | N | Rho | p-value |
| --- | --- | --- | --- |
| $A_H$>$G_H$ (Spearman) | 19 | 0.4387708 | 0.0602 |
| $A_H$>$G_H$ (Pearson) | 19 | 0.8239473 | 1.448e-05 |

Oxygen consumption by standard metabolism was retrieved from FishBase database (https://www.fishbase.se/home.htm).
